# Neuro-mimetic reprogramming and epigenetic unlocking drive lung cancer stem cell adaptation to the brain microenvironment

**DOI:** 10.64898/2026.10.29.762751

**Authors:** Elisa Arias-Romero, Ángeles Fernández-Fondevila, Alejandra Macías-Cerro, Beatriz Fernández-Roldán, Pilar Cerveró-García, Vinothini Rajeeve, Rafael Coveñas, Arantxa Tabernero, Pedro R. Cutillas, Maruan Hijazi

**Affiliations:** Group GIR BMD (Bases Moleculares del Desarrollo), University of Salamanca, Salamanca, Spain; Institute for Neuroscience of Castilla y León (INCyL), iBRAINS-IN-CYL, Salamanca, Spain; Institute of Biomedical Research of Salamanca (IBSAL), Salamanca, Spain; Barts Cancer Institute, Queen Mary University of London, Charterhouse Square, London, EC1M 6BQ, United Kingdom; Institute of Parasitology and Biomedicine “López-Neyra” (IPBLN-CSIC), Granada, Spain

**Author notes:** Correspondence (M.H).

**Keywords:** brain metastasis, lung cancer stem cells, phosphoproteomics, phenotypic plasticity, neural mimicry

## Abstract

Brain metastasis represents a lethal complication of non-small cell lung cancer, driven by the ability of disseminated tumor cells to colonize the central nervous system microenvironment. Although cancer stem cells (CSCs) are recognized as key drivers of metastatic initiation, whether their contact with the brain parenchyma induces terminal differentiation or adaptive plasticity remains controversial. Here, using an in vivo model of lung-derived CSCs injected intracranially, combined with quantitative total proteomics and phosphoproteomics, we demonstrate that contact with the brain niche triggers a neuro-mimetic reprogramming rather than terminal differentiation. Lung CSCs desensitize their systemic metastatic program while epigenetically unlocking neural stem cell networks, characterized by the upregulation of Ngfr, Fabp7, Pbx3, Vgf, and S100b, concurrent with the downregulation of the DNA methyltransferase Dnmt3a and the cytosolic exclusion of Id1. Phosphoproteomic profiling revealed post-translational adaptations, highlighted by hyperphosphorylation of Nestin at S957 and S963. Furthermore, kinase substrate enrichment analysis uncovered high activity of Prkcq (PKCθ) and Map3k7 (TAK1), shifting the cellular state toward structural remodelling and microenvironmental stress tolerance. Functional validation assays using selective inhibitors confirmed that these kinase networks represent adaptive liabilities within tissue constraints rather than basic in vitro survival requirements. Together, our findings demonstrate that lung CSCs exploit a phenotypic plasticity to adapt to the brain niche, acquiring a neural-glial disguise that supports vascular co-option, immune evasion, and metastatic colonization.

## Introduction

Non-small cell lung cancer constitutes the leading cause of cancer-related mortality worldwide, with brain metastasis representing its most devastating complication, affecting up to 40% of patients during the disease course [1,2]. Despite recent therapeutic advances incorporating targeted therapies and immune checkpoint blockade, the prognosis for patients with central nervous system (CNS) dissemination remains remarkably poor [3,4]. This clinical resistance stems from the unique biological architecture of the brain parenchyma, which acts as a niche protected by the blood-brain barrier and a highly specialized extracellular matrix, imposing unique biophysical and immunological pressures on invading carcinoma cells [5,6].

Within the intratumoral hierarchy, the subpopulation of cancer stem cells (CSCs) has been identified as the primary engine driving tumor initiation, therapeutic resistance, and distant metastatic seeding [7,8]. During the metastatic cascade, lung CSCs must intravasate from the primary lung tissue, survive in systemic circulation, and cross the cerebral microvasculature to establish parenchymal micrometastases [9]. However, the phenotypic fate of lung-derived CSCs upon initial contact with the brain microenvironment remains a matter of intense debate. While historical paradigms postulated that the brain stroma might induce differentiation in exogenous carcinoma cells as a host defence mechanism, emerging evidence indicates its capacity for phenotypic plasticity, where metastatic cells mimic structural and functional traits of native neural and glial lineages, a phenomenon termed neural or glial mimicry [10].

Executing this neuro-mimetic adaptation requires metastatic cells to actively reconfigure their transcriptional machinery and cytoskeletal framework [11]. To colonize the brain parenchyma, lung CSCs attenuate their active systemic mesenchymal invasion program and engage epigenetic unlocking pathways that allow the expression of neural progenitor factors. Cell-surface markers such as the nerve growth factor receptor Ngfr (CD271) and lipid chaperones like Fabp7 serve as pivotal regulators of self-renewal and brain tropism [12,13], whereas homeobox transcription factors such as Pbx3 coordinate resistance against differentiation [14]. This identity switch relies on the downregulation of repressive DNA methyltransferases, such as Dnmt3a, and chromatin corepressor complexes [15,16], liberating the cell from the lineage constraints of its tissue of origin.

Parallel to genomic reprogramming, physical colonization of the brain parenchyma entails a mechanical remodelling of the cytoskeleton and extracellular matrix [17,18]. Disseminated tumor cells abandon destructive peripheral invasion mechanisms in favour of vascular co-option, sliding along the outer surface of pre-existing microvessels via protective serpins like Serpine 2 and specialized basement membrane components [17,19]. Throughout this morphological transition, intermediate filament proteins such as Nestin undergo dynamic spatial reorganization regulated by post-translational phosphorylation events that alter filament assembly [20]. Moreover, interactions with resident neuro-immune cells (microglia and reactive astrocytes) force metastatic cells to deploy immune evasion strategies, combining the downregulation of major histocompatibility complex class I (MHC-I) antigen presentation machinery with the expression of myeloid cell surface markers [21,22].

To rapidly coordinate these multifaceted adaptive responses against physical and metabolic stress in the neural niche, CSCs rely on the hyperactivation of specific kinase networks [23]. Signaling cascades mediated by kinases such as Prkcq (PKCθ) and Map3k7 (TAK1) emerge as central hubs controlling actin dynamics, intercellular junctions, and resistance to microenvironmental inflammation [24,25]. Consequently, quantitative profiling of the proteome, phosphoproteome and kinase activity in in vivo models is essential to uncover how lung CSCs adapt their phenotype to the brain microenvironment and to identify vulnerabilities that can be exploited therapeutically.

## Results

### Spatial reorganization of the Nestin cytoskeleton and subcellular redistribution of the transcriptional regulator Id1

Firstly, we assessed the effect of the brain microenvironment on injected lung CSCs (3LL stem subpopulation) using an in vivo model of lung cancer brain metastasis. To evaluate whether the phenotypic changes induced by the brain microenvironment involved variations in classical markers of differentiation and stemness plasticity [26,20], we analyzed the expression levels and subcellular localization of Nestin and Id1 via Western Blot and immunocytochemistry (ICC) in 3LL pre-implanted (named as 3LL preC) and 3LL post-implanted (named as 3LL postC and isolated after establishing a brain tumor for 24 days) cells (Figure 1). Western blotting analysis revealed no statistically significant differences in total Nestin protein abundance between pre-implanted and post-implanted conditions (Figure 1A). However, ICC analysis demonstrated a shift in the spatial compartmentalization of Nestin. While 3LL preC cells exhibited a highly condensed, perinuclear Nestin signal displaying marked spatial overlap with DAPI nuclear staining, 3LL postC cells displayed a diffuse, filamentous network extending throughout the entire cytoplasm (Figure 1B, upper panel). This finding demonstrates a structural reorganization of the intermediate filament network in response to contact with the brain niche [27].

**Figure 1.**
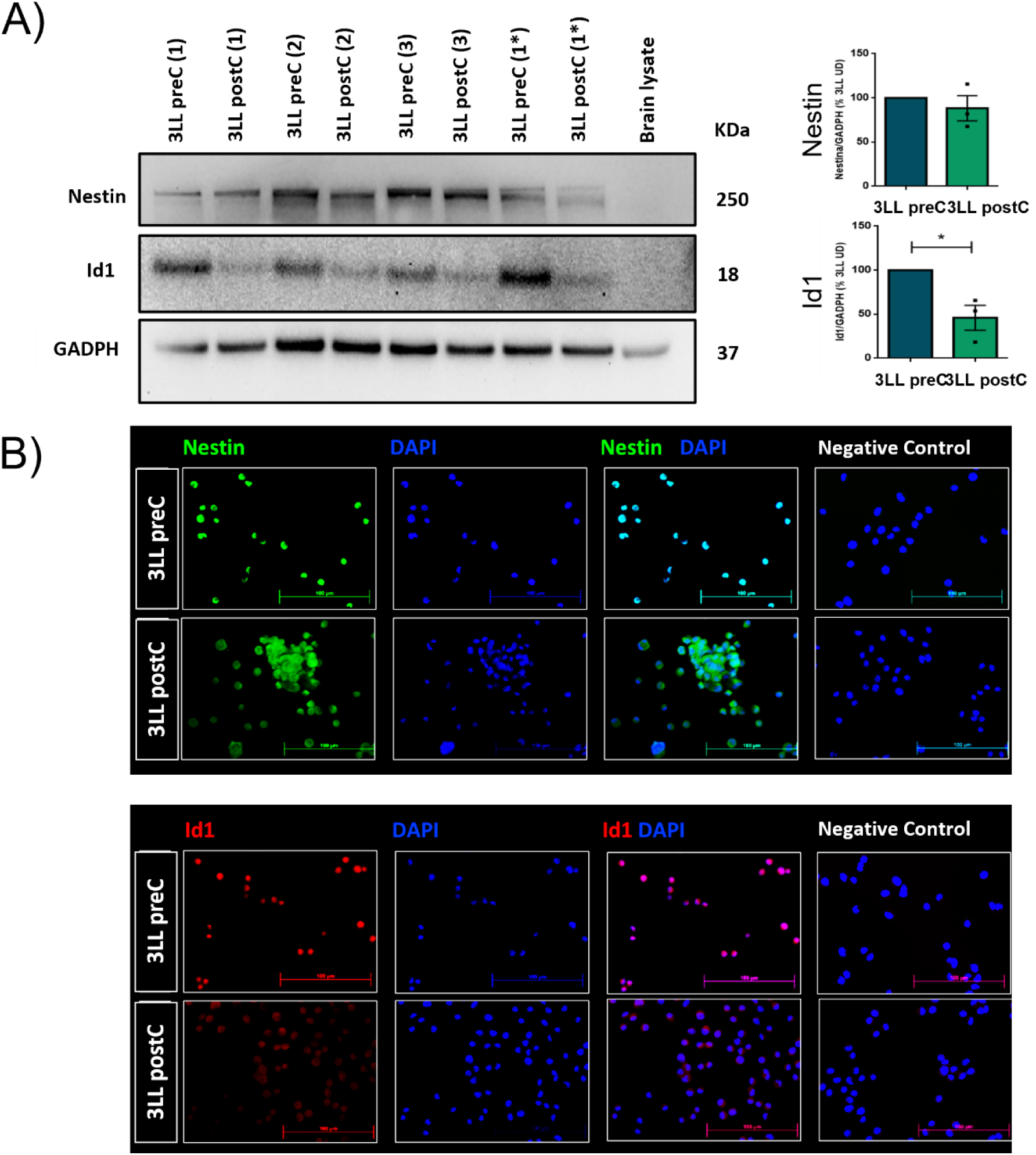
Effect of the brain microenvironment on differentiation markers and stemness plasticity. A) 3LL postC denoted 3LL stem cells post brain implantation (isolated after establishing a brain tumor for 24 days) meanwhile numbers 1, 2 and 3 denoted biological replicates. Biological replicate 1 (1*) was loaded twice in the gel. Values were normalized to GAPDH levels and are expressed as ± SEM; n=3. *p<0.05. B) Representative images of 3LL stem cells (3LL preC) and isolated 3LL stem cells post brain implantation (3LL postC). Immunocytochemistry stained with Nestin (green), Id1 (red) and DAPI (blue) showed subcellular redistribution for both Nestin and Id1 in 3LL postC. Scale bar: 100 µm.

In contrast to Nestin, total Id1 protein expression experienced a significant downregulation in 3LL postC cells compared to the 3LL preC control (Figure 1A). At the subcellular level, 3LL preC cells displayed an exclusively nuclear localization of Id1, co-localizing strongly with DAPI chromatin staining [28]. Conversely, in 3LL postC cells, in addition to a marked decrease in overall fluorescence intensity, the remaining Id1 signal was restricted to the cytosolic compartment (Figure 1B, lower panel). This loss of nuclear localization confirms the functional suppression of Id1-mediated transcriptional repression over nuclear bHLH factors in the 3LL postC model [28,26].

### Lung CSCs undergo neuro-mimetic reprogramming and reinforcement of stemness

To further elucidate the molecular basis of these findings, we performed a mass spectrometry (MS)-based quantitative proteomics analysis. The experiment compared our two conditions (3LL postC vs 3LL preC) using two biological and two technical replicates across eight LC-MS/MS runs. This approach identified over 50,000 peptides corresponding to nearly 4,500 protein groups. Among them, 43 and 56 proteins were found to be significantly upregulated and downregulated, respectively (Supplementary Figure 1A). Specifically, the top 20 statistically significant differentially expressed proteins (p < 0.05) were ranked according to their fold change (Supplementary Figure 1B). Comparative quantitative proteomic profiling between lung CSCs exposed to the brain microenvironment in vivo (3LL postC) and control cells maintained in vitro (3LL preC) revealed a slightly shift in cellular protein abundance. Firstly, we observed a significant upregulation of self-renewal markers and CNS progenitor lineage factors in postC cells, including the neurotrophin receptor Ngfr (a driver of stemness and perivascular invasion) [29] and the brain fatty acid-binding protein Fabp7 (a marker of radial glia and neural progenitor niche) (Figure 2A) [13]. Furthermore, the differentiation-blocking transcription factor Pbx3 [14], the neuropeptide Vgf [30] and the glial lineage marker S100b displayed statistically significant increases (Figure 2A, upper panel). In addition, we detected a significant decrease in Sufu [31], a key negative regulator of the oncogenic Hedgehog pathway, and Dpysl3 [32], a mediator of neuronal differentiation (Figure 2A, lower panel). Thus, proteomic analysis confirmed the induction of neural progenitor markers and the concomitant repression of neuronal differentiation drivers.

**Figure 2.**
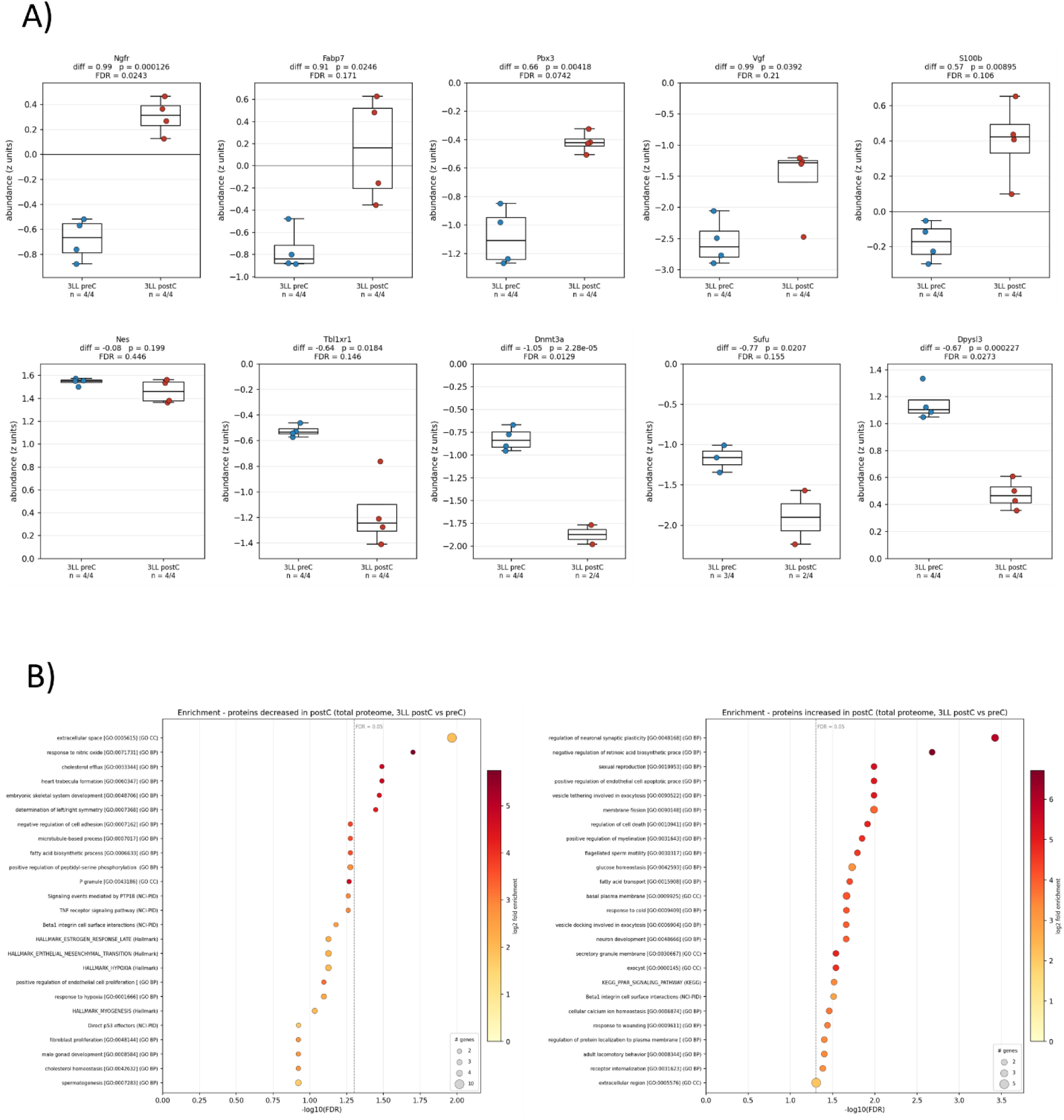
Brain microenvironment contact induces neuro-mimetic reprogramming, stemness reinforcement and functional pathway rewiring in lung cancer stem cells. A) Relative protein abundance of key functional markers comparing control 3LL preC and intracranially adapted 3LL postC cells, determined by quantitative mass spectrometry (n=4 replicates per group). Boxplots display median, interquartile range and individual data points. Statistical significance, mean abundance difference, p-values (Welch’s test) and FDR are indicated above each plot. B) Functional pathway enrichment analysis based on total proteome alterations. Left and and right panels denoted enriched terms among significantly decreased or increased, respectively, proteins in postC cells. Dot size corresponds to the number of identified genes per term (n), dot colour represents the log2 fold enrichment scale and the x-axis shows the significance threshold (FDR = 0.05) indicated by the dashed line.

Furthermore, the proteomic profile demonstrated a remodelling of chromatin-modifying machinery and cellular plasticity factors. Notably, we detected a reduction in the de novo DNA methyltransferase Dnmt3a [15] and Tbl1xr1, an essential subunit of the N-CoR corepressor complex [16] (Figure 2A, lower panel). Moreover, exposure to the brain microenvironment induced a restructuring of stromal interaction factors and membrane architecture [17]. Specifically, postC cells significantly downregulated the laminin subunit Lamb1, the vascular cadherin Cdh13, the actin organization factor Pfn2, and the caveolar marker Cav1 (Supplementary Figure 2A). Conversely, postC cells exhibited increased levels of the cell-communicating tetraspanin Cd81, the glial serpin Serpine 2, and the basement membrane constituent Col18a1 (Supplementary Figure 2B). Collectively, these shifts indicate an adaptive reorganization of extracellular matrix interactions, cell adhesion, and vascular niche alignment [17,19]. Finally, proteomic characterization revealed alterations in immune presentation and myeloid-like markers. We observed a significant reduction in the major histocompatibility complex class I molecule H2-K1 and the vascular cell adhesion molecule Vcam1 (Supplementary Figure 2A). Conversely, postC cells upregulated the lysosomal enzyme Hexb, the immune modulator Aif1l, and the oxidative stress/immune sensor Keap1, pointing toward a modulation of immune interaction mediators in the neural niche (Supplementary Figure 2B) [21,22].

Then, we conducted pathway enrichment analysis using proteins significantly upregulated following intracranial contact. Remarkably, the top-ranked pathways strongly pointed to the acquisition of a neural lineage phenotype (Figure 2B, right panel). The most prominent terms included “regulation of neuronal synaptic plasticity,” “neuron development,” and “positive regulation of myelination”. Additionally, we identified significant enrichment in “fatty acid transport” and “KEGG PPAR signaling pathway”. S100b and Ngfr directly contribute to terms associated with neuronal development and synaptic plasticity, whereas the elevated expression of Fabp7 aligns precisely with enriched fatty acid transport and PPAR signaling [13]. Furthermore, enrichment in synaptic plasticity pathways matched with the suppression of classical mesenchymal invasion programs, evidenced by a significant reduction in “Hallmark epithelial mesenchymal transition” [9]. The downregulations of Cav1 and Dpysl3 contribute to slow down peripheral mesenchymal migration and disassembling the initial cytoskeleton [32], a permissive transition enabled by the loss of the epigenetic repressor Dnmt3a [15]. Regarding more downregulated pathways, the significantly decreased terms included “extracellular space,” “negative regulation of cell adhesion,” and “Beta1 integrin cell surface interactions” (Figure 2B, left panel), confirming the deconstruction of primary pulmonary stromal adhesion networks. This loss of adhesion could be partly mediated by the reduction of Lamb1, Cdh13, and Pfn2 (Supplementary Figure 2A). Finally, we observed a decrease in the “TNF receptor signaling pathway”. Because TNF signaling serves as a primary driver of inflammatory responses and leukocyte recruitment [21,22], its downregulation matches with the reduction of Vcam1 and contributes to the reduction of H2-K1 (MHC-I) (Supplementary Figure 2A), thereby restricting tumor antigen presentation. Simultaneously, postC cells upregulate survival factors against immune stress, such as Keap1, and emulate lysosomal myeloid components, including Hexb and Aif1l, to facilitate immune tolerance in the brain parenchyma (Supplementary Figure 2B) [22].

### Driving role of cancer stem cells on the brain microenvironment by phosphoproteomics

To better understand the signaling pathways involved in CSCs phenotypic plasticity, we performed a MS-based phosphoproteomics experiment that identified more than 12,000 phosphopeptides (Figure 3). A total of 2,557 proteins were identified, of which 2,268 (88.7%) were phosphorylated. Comparing 3LL preC vs 3LL postC conditions, the volcano plot demonstrated that physical contact with the brain microenvironment exerted a substantial impact on the phosphoproteome of post-implanted cells (519 increased and 577 decreased phosphosites) (Figure 3A). Moreover, the top 20 statistically significant differentially expressed proteins (p < 0.05) were ranked according to their fold-change (Figure 3B). Analysis of the phosphoproteomic dataset identified site-specific phosphorylation events on key functional proteins. We detected a significant increase in Nestin phosphorylation at S957 and S963 in 3LL postC cells, modifications that could induce filament disassembly and intermediate filament network relaxation (Supplementary Figure 3A). This result indicates that the brain microenvironment activates specific kinases that hyperphosphorylate Nestin, promoting cytoskeletal plasticity [27]. This structural transition provides tumor cells with the biophysical flexibility required to deform, migrate, and infiltrate the brain parenchyma [18].

**Figure 3.**
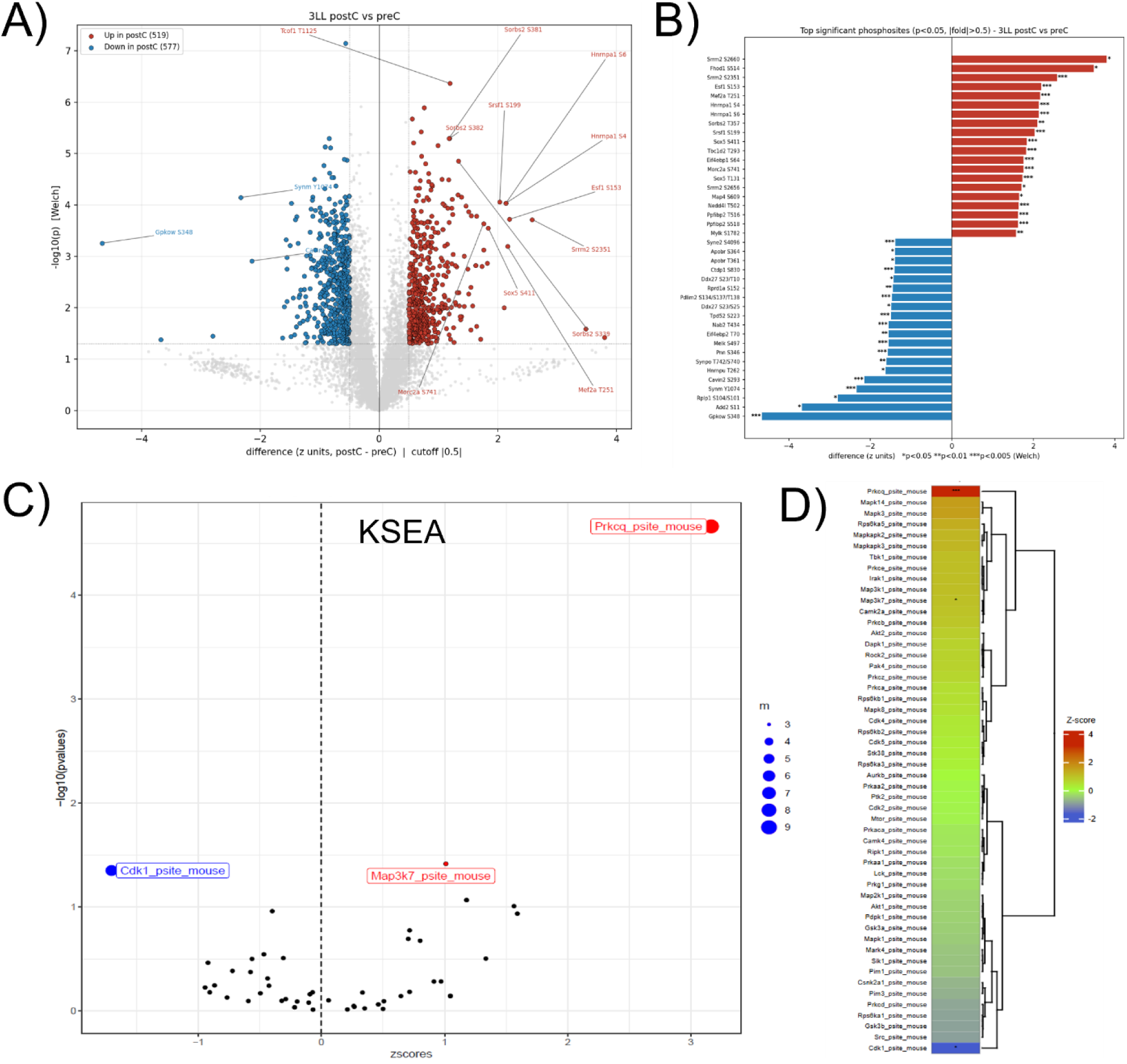
A) Volcano plot showing phosphosites significantly decreased (blue) and increased (red) in 3LL postC compared to 3LL preC (control). Phosphosites were identified and quantified using MS-based phosphoproteomics (p-value threshold was 0.01). B) Top 20 increased or decreased phosphosites were labelled, p < 0.05). C) Volcano plot showing kinase activities increased (red) or decreased (blue) in 3LL postC compared to control 3LL preC (quantified using kinase substrate enrichment analysis or KSEA). Labels indicate significant kinase activities. D) Quantification from changes in substrate phosphorylation calculated by KSEA ranked a panel of kinases associated to the phosphosites detected in the analysis. Those sites inferred kinase activities relative to control 3LL preC and showed as z-score.

We also observed a significant increase in Ngfr phosphorylation at S314, indicating structural modification of its intracellular domain that could promote active downstream signaling toward NF-κB survival pathways or RhoA-mediated motility (Supplementary Figure 3A) [29]. Thus, 3LL postC cells could use microenvironmental trophic signals to drive phenotypic plasticity [10]. Conversely, Dpysl3 showed a significant decrease in phosphorylation at T509 and S522 in 3LL postC cells (Supplementary Figure 3A). Dpysl3 regulates microtubule dynamics and growth cone collapse and its phosphorylation by kinases such as Gsk3β or Cdk5 inhibits tubulin binding and induces cytoskeletal collapse [32]. The dephosphorylated Dpysl3 form enriched in our dataset could be active in stabilizing microtubule dynamics, facilitating filopodia formation and neuro-glial invasive mimicry, a mechanism required for vascular co-option [17]. Thus, the phosphoproteome demonstrates that contact with the brain microenvironment not only erases original lineage constraints epigenetically [15], but triggers post-translational responses via Nestin, Ngfr, and Dpysl3 phosphorylation to direct cytoskeletal restructuring [11,32], enabling tumor cells to physically adapt to the neural niche.

Phosphoproteome-based GO enrichment analysis confirmed that the most robust upregulation occurred in structural and membrane-associated pathways, including “cytoskeleton”, “adherens junction”, and “plasma membrane” in agreement with Nestin phosphorylation dynamics (Supplementary Figure 3B, right panel) [27]. This reorganization grants cells the biomechanical flexibility needed for parenchymal infiltration and vascular co-option [17]. Conversely, the most significantly downregulated phosphoproteomic terms involved “transcription corepressor activity”, in agreement with the functional inactivation of Id1 and Dnmt3a corepressor networks (Supplementary Figure 3B, left panel). Finally, we detected a decreased enrichment in “KEGG antigen processing and presentation”, confirming that antigen presentation machinery could be disassembled at the post-translational level to evade cytotoxic T lymphocyte recognition in the brain [21,22].

To estimate changes in kinase activity based on collective substrate phosphorylation dynamics, we performed kinase substrate enrichment analysis (KSEA) (Figure 3C) [23]. KSEA uncovered a statistically significant increase in the activity of Prkcq (PKCθ) and Map3k7 (TAK1) in 3LL postC cells. Prkcq is a major modulator of actin cytoskeleton dynamics and adherens junctions [24], providing a functional link to the hyperphosphorylation of structural proteins and the spatial redistribution of Nestin filaments [27]. Meanwhile, Map3k7 serves as a central signaling node coordinating cellular stress, survival, and inflammatory pathways (including NF-κB and p38 MAPK) [25]. Its activation in post-implanted cells might be crucial to resist microenvironmental stress, driving the attenuation of immune presentation while supporting neuro-mimetic transcriptional reprogramming [21,25]. Other kinase activities were found and listed in Figure 3D but not labelled in 3C since they do not reach significance threshold.

### Cell survival analysis as a functional validation to exploit cancer stem cells vulnerabilities

To functionally validate the high activity of PKCθ and TAK1 obtained by KSEA, we performed cell survival assays using selective pharmacological inhibitors (Figure 4). For PKCθ, we tested three distinct compounds: Sotrastaurin (pan-PKC inhibitor), Darovasertib (PKCα and PKCθ inhibitor), and a PKCθ isoform-selective inhibitor named PKC theta inhibitor. Cell viability exhibited a stronger decreasing trend in 3LL postC cells, consistent with elevated PKCθ activity. However, these differences did not achieve statistical significance under culture conditions. Similarly, TAK1 inhibition using Takinib produced no significant viability differences between preC and postC cells. These findings indicate that cell survival assays do not fully capture the role of kinases whose primary functions are adaptive, structural, and microenvironment-dependent (such as cytoskeletal remodelling, stromal interaction, and immune evasion) rather than intrinsic survival.

**Figure 4.**
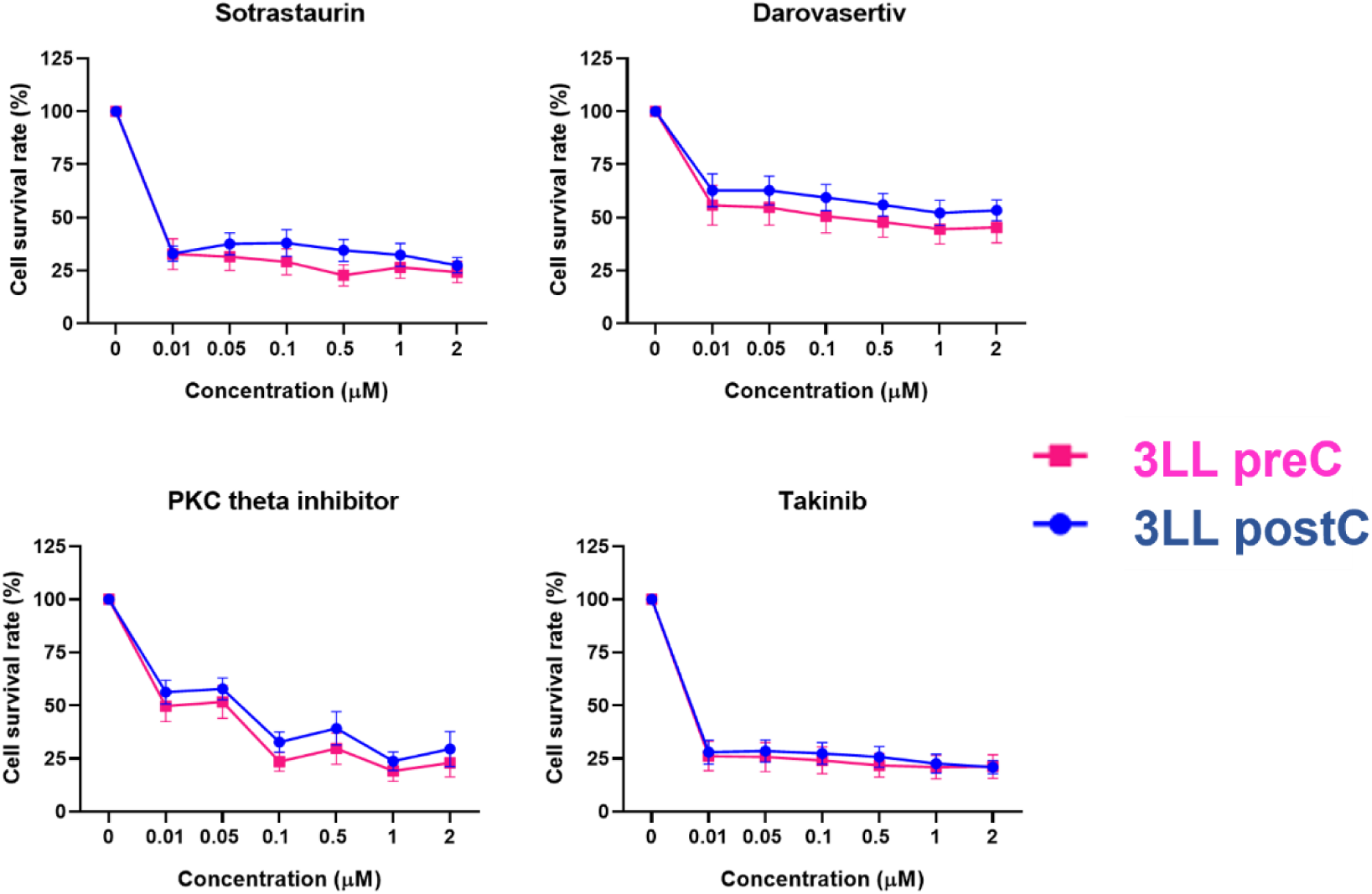
Pharmacological validation of PKC theta and TAK1 kinase inhibitors in 3LL cancer stem cells. Dose-response cell survival curves comparing control 3LL preC and brain-adapted cells (3LL postC) treated for 48 hours with increasing concentrations of specific kinase inhibitors: sotrastaurin (top left), darovasertib (top right), PKC theta inhibitor (bottom left) and the selective TAK1 inhibitor takinib (bottom right). Cells were incubated with 5 % resazurin for 4 hours, and fluorescence was measured at λ_ex_ = 565 nm and λ_em_ = 590 nm (mean and SEM, n=3 biological replicates).

## Discussion

Metastatic colonization of the brain parenchyma by lung cancer stem cells represents an extraordinary adaptive challenge [1,2]. Upon intracranial transplantation into a microenvironment fundamentally foreign to their primary pulmonary niche, our proteomic and phosphoproteomic findings demonstrate that lung CSCs do not undergo differentiation, but rather perform a neuro-mimetic reprogramming while reinforcing their stemness state [10,12].

Contact with the brain parenchyma induces the expression of self-renewal and tissue plasticity effectors such as Ngfr and Fabp7, key regulators of radial glia progenitors and CSC survival [12,13]. This transition is reinforced by the increase of the developmental factor Pbx3 [14] and the neuropeptide Vgf [30], paired with the downregulation of Sufu, a direct suppressor of Hedgehog signaling. Sufu downregulation triggers oncogenic Hedgehog activation [31], whereas the decrease in Dpysl3 confirms the repression of neuronal differentiation programs [32]. At the chromatin level, this switch is supported by the loss of the DNA methyltransferase Dnmt3a [15] and the NCoR corepressor subunit Tbl1xr1 [16]. Dnmt3a attenuation promotes the demethylation of genomic regions previously silenced in the pulmonary lineage [15], allowing tumor cells to respond flexibly to signals from the brain microenvironment [10].

Physical adaptation to the vascularized brain tissue [17,18] is evidenced by the downregulation of peripheral adhesion markers and the activation of vascular support networks. The decrease in Lamb1 and Cdh13, alongside reduced Cav1 and Pfn2 levels, indicates the disassembly of invasion machinery [9]. In its place, elevated Serpine 2 and Col18a1 levels support vascular co-option and matrix preservation [17,19], complemented by increased Cd81, a tetraspanin important for vesicular packaging and trophic glia interaction. Furthermore, crosstalk with resident neuro-immune cells reveals an adaptive strategy [21,22]. On one hand, downregulation of H2-K1 and Vcam1 impairs antigen presentation, providing a classic mechanism to evade CD8+ cytotoxic T lymphocyte surveillance [21]. On the other hand, upregulation of Aif1l and Hexb points to a phenomenon of myeloid/microglial mimicry [22], where tumor cells could adopt molecular features of resident microglia to escape immune recognition.

Phosphoproteomic profiling confirmed that this adaptive phenotype is orchestrated by post-translational modifications [23]. Hyperphosphorylation of cytoskeletal and adherens junction components [27] drives the spatial reorganization of the Nestin filament network, increasing biomechanical flexibility [33]. Moreover, contact with the neural niche induces the dephosphorylation and functional inactivation of transcriptional corepressors and histone modifiers [16,26]. This, combined with decreased antigen presentation pathways [21], could allow tumor cells to arrange a neuro-mimetic program while remaining immunologically silent [10,21]. KSEA provided a systemic view of the kinase signaling driving this transition [23] and demonstrated a significant increase in the activity of Prkcq (PKCθ) and Map3k7 (TAK1) [24,25], together with a marked reduction in Cdk1 activity. Downregulation of Cdk1, a master regulator of cell cycle progression, highlights that tumor cells temporarily decelerate cell cycle proliferation to assign energetic resources toward kinase networks leading migration, cytoskeletal remodelling, and microenvironmental stress survival. This kinase rewiring confirms that brain colonization relies on structural and invasive adaptation rather than uncontrolled cell division [17]. For further detail, individual substrates that infer kinase activity for each of the three kinases are highlighted in Supplementary Figure S4.

To functionally validate the elevated activity of PKCθ and TAK1, we tested cell sensitivity to selective pharmacological inhibition. Inhibition of PKCθ across three independent compounds (Sotrastaurin, Darovasertib, and the selective PKC theta inhibitor) revealed a consistent trend toward reduced viability in 3LL postC cells, although these differences did not reach statistical significance in suspension cultures. In addition, TAK1 inhibition using Takinib produced no differential effect. These results indicate that hyperactivated PKCθ and TAK1 do not primarily sustain basic cell-autonomous proliferation in vitro, but function as adaptive signaling nodes [24,25]. Within the brain parenchyma, these kinases coordinate biomechanical plasticity, cytoskeletal reorganization, and local immune evasion [24,25]. Consequently, their functional dependency becomes critical under microenvironmental stress and tissue invasion rather than during cell culture.

## Supplementary Figures

**Figure S1.**
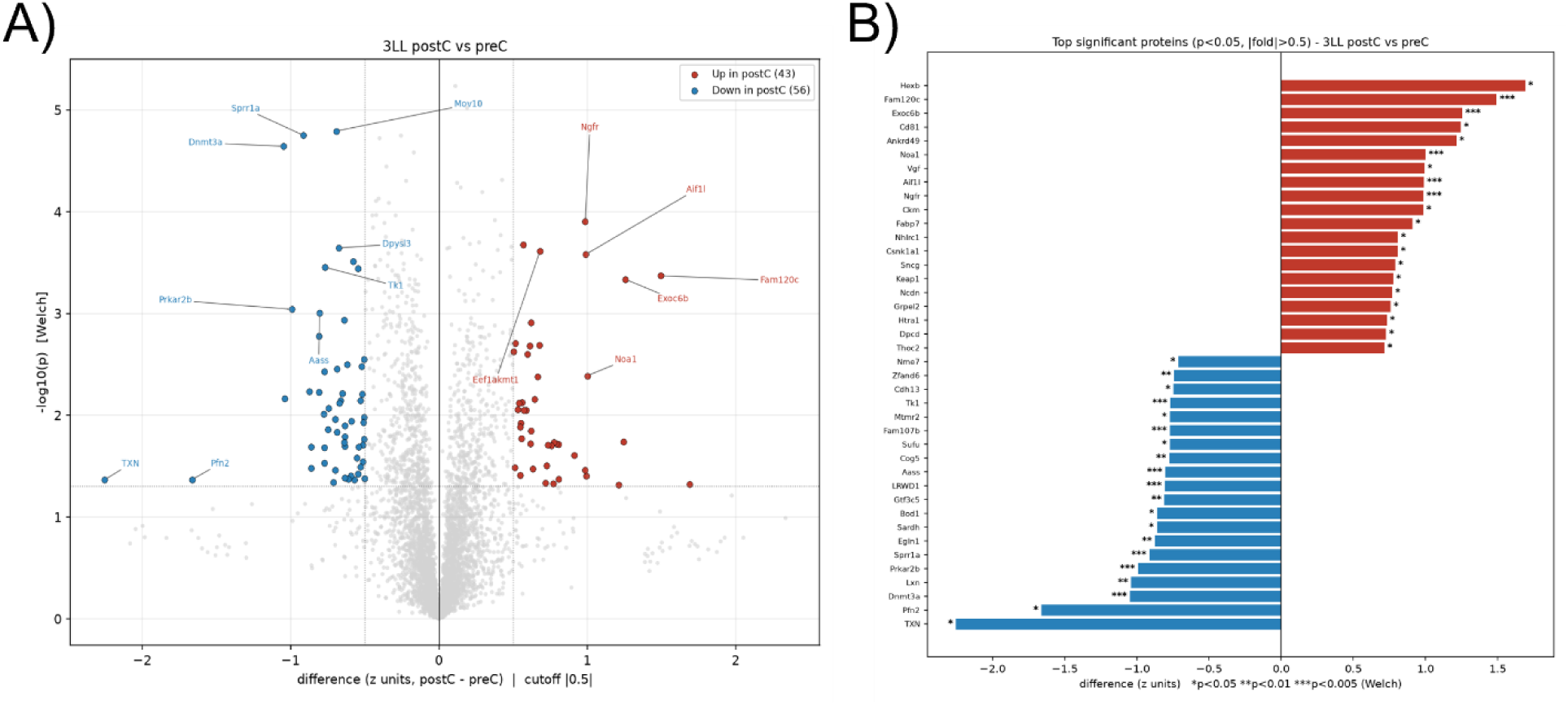
Quantitative proteomic analysis of 3LL lung cancer stem cells following intracranial adaptation. A) Volcano plot showing differential protein expression between 3LL postC (intracranially injected cells) and control cells (3LL preC). Statistically significant upregulated proteins in 3LL postC are highlighted in red, whereas downregulated proteins are shown in blue. B) Top statistically differentially expressed proteins, ranked by fold change (p < 0.05). Statistical significance was evaluated using Welch’s test (*p < 0.05, **p < 0.01, ***p < 0.005).

**Figure S2.**
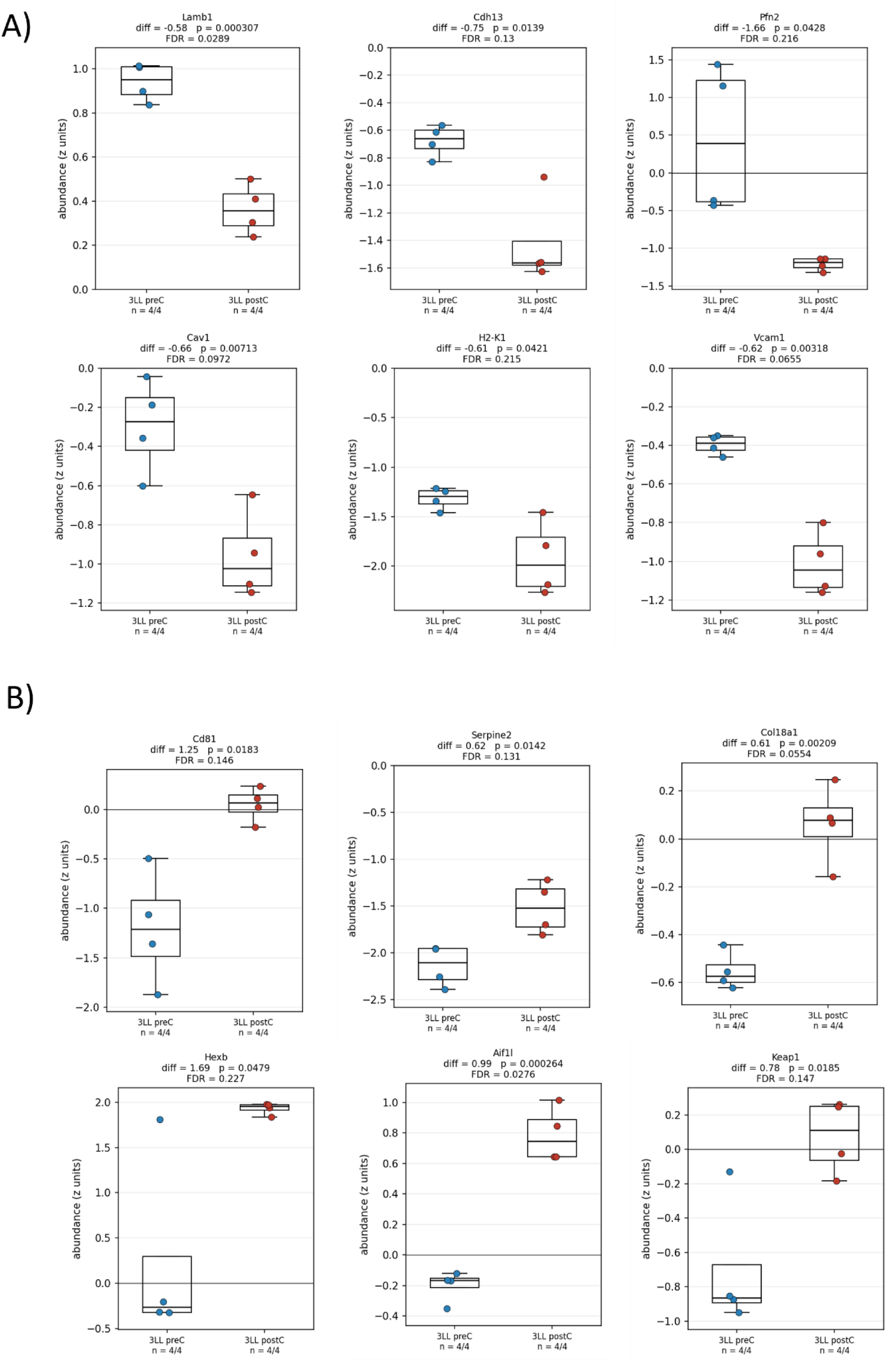
Quantitative proteomic alterations in stromal adhesion, vascular co-option and immune modulation factors upon intracranial adaptation. A) Relative protein abundance of significantly decreased factors comparing control 3LL preC (blue dots) and 3LL postC (red dots), determined by mass spectrometry (n = 4 replicates per group). B) Relative protein abundance of significantly increased factors in 3LL postC cells. Boxplots present median values, interquartile ranges and individual data points. Mean abundance differences, unadjusted p-values (Welch’s test) and FDR are specified above each boxplot.

**Figure S3.**
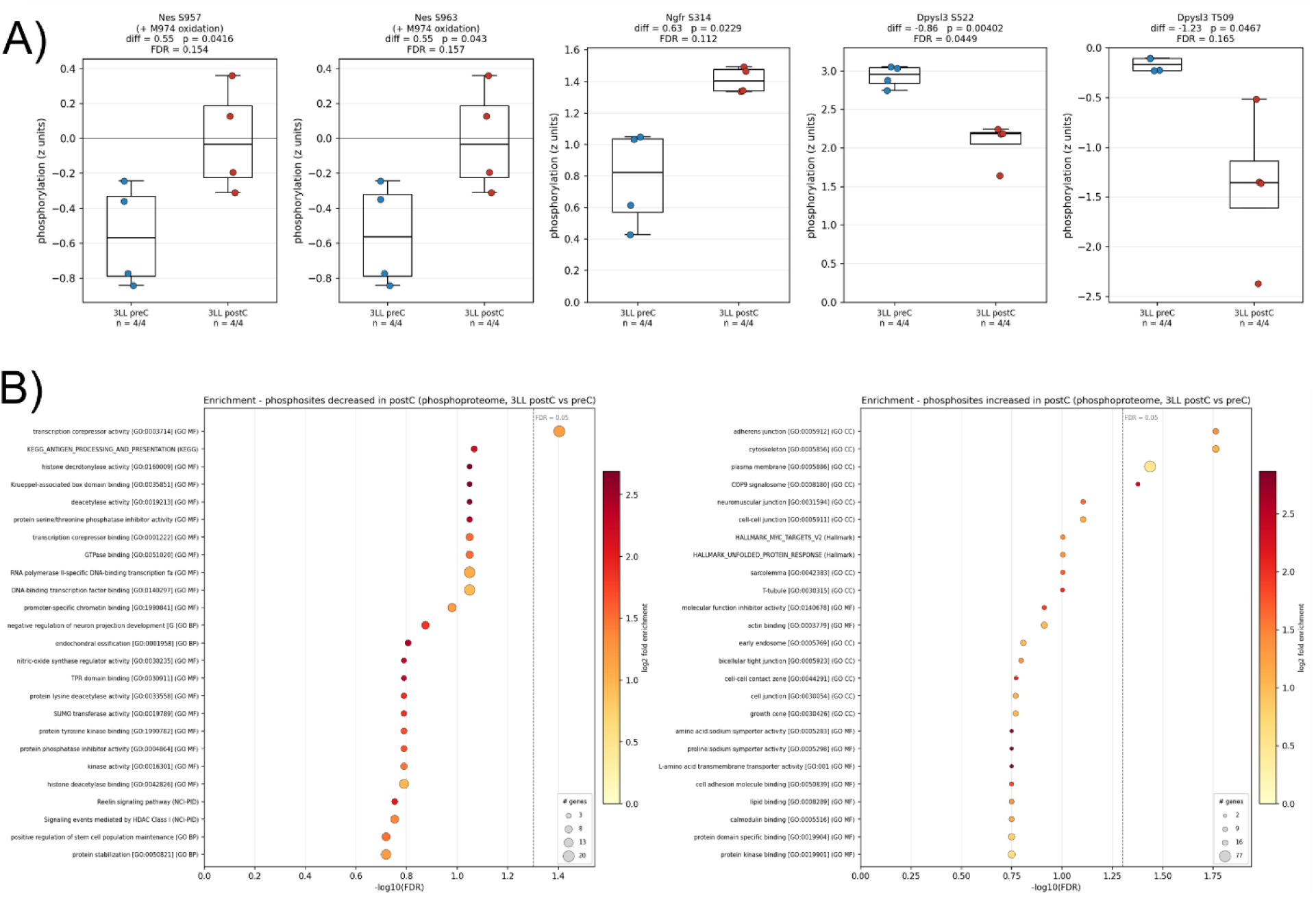
Phosphoproteomic profiling and pathway enrichment analysis in intracranial adapted lung cancer stem cells. A) Quantitative phosphorylation levels of key functional proteins comparing control 3LL preC (blue dots) and 3LL postC cells (red dots), determined by MS-based phosphoproteomics (n = 4 replicates per group). Boxplots display median, interquartile ranges and individual data points. Mean phosphorylation differences, p-values (Welch’s test) and FDR are indicated above each boxplot. B) Phosphoproteome-based pathway enrichment analysis. Right and left panels refer to enriched pathways among phosphosites significantly increased or decreased, respectively, in postC cells. Dot size corresponds to the number of identified genes per term (n), dot colour represents the log2 fold enrichment scale and the x-axis shows the significance threshold (FDR = 0.05) denoted by the dashed vertical line.

**Figure S4.**
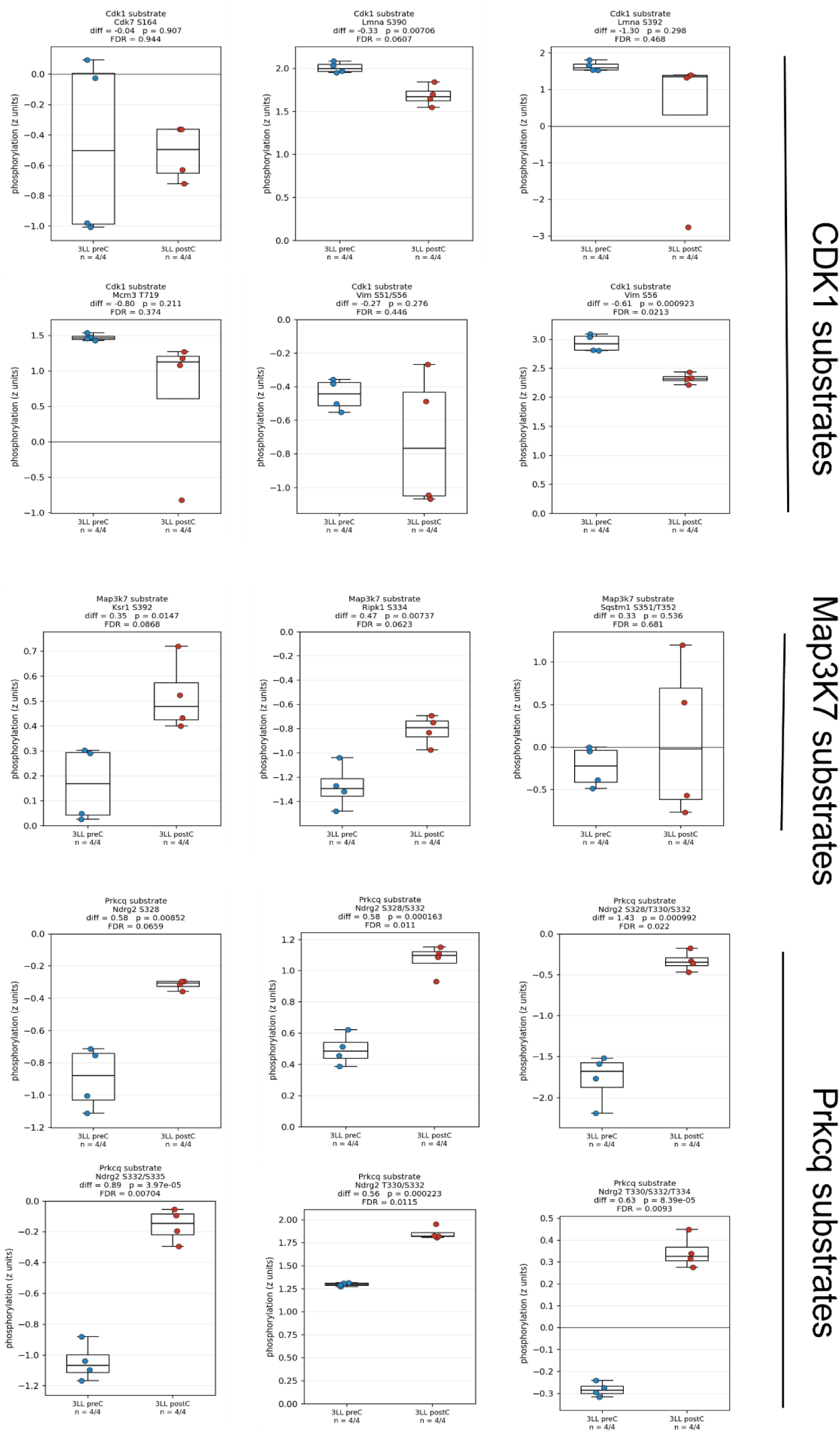
Quantitative phosphorylation of kinase substrates inferring Cdk1, Map3K7 and Prkcq activity changes upon intracranial adaptation. Phosphorylation levels quantified by mass spectrometry of specific phosphosites corresponding to validated substrates for the named kinases, comparing control 3LL preC (blue dots) and 3LL postC (red dots) (n= 4 replicates per group). Boxplots depict median values, interquartile ranges and individual data points. Mean phosphorylation differences, p-values (Welch’s t-test) and FDR are indicated above each plot.

## Material and Methods

### Animals

C57BL/6 mice were obtained from Charles River Laboratories in agreement with the Experimental Animal Service of the University of Salamanca. All animal procedures were previously approved by the research ethics committee of the University of Salamanca (CBE 697 and 838) and the Junta de Castilla y León approval, following the European Community Council directives (2010/63/UE) and the Spanish law RD 53/2013 for the use and care of laboratory animals.

### Cells

Mouse lung cancer Lewis Lung Carcinoma (3LL, ATCC Ref CRL-1642) cells were cultured in non-adherent conditions (3LL cancer stem cells, 3LL-CSCs). Briefly, adherent 3LL cells were cultured in DMEM supplemented with progressively lower FBS concentration, starting at 10%. After 3 days, FBS concentration was reduced to 5%, and 3 days later to 2.5%, following the methodology described by Yi et al [34]. Then, after 3-5 days, when cells started to detach from the plate surface, they were transferred to non-treated T25 flasks with stem cell medium DMEM/F-12 with glutamine supplemented with 1.45% glucose, 1% MEM non-essential amino acids 100x, 1% Pen/Strep, 0.16 % BSA, 0.2% β-mercaptoethanol and supplemented with 1% B27, 20 ng/mL human EGF and 20 ng/mL human b-FGF. Non-adherent cultures (3LL-CSCs, also referred to as 3LL preC) were considered established after 3-4 weeks culture in stem cell medium.

### Stereotactic surgery for intracranial 3LL implantation

3LL-CSCs were intracranially implanted into the brains of 8-week-old C57BL/6 mice as previously described [35]. Mice were anaesthetized using isoflurane while placed in a stereotaxic frame. Stereotactic coordinates were used (1.5 mm AP + 1.5 mm LM to bregma) to trephine the skull and make the brain tissue accessible for injection. 1uL saline containing 5×10^3^ cells was injected at -1mm deep into the brain tissue using a Hamilton microsyringe coupled to a 26G needle.

### Stem cells isolation from mice brains

Isolated cells will be referred to as post-brain implant 3LL (3LL postC). Briefly, the brains were thawed and the tumors then cut, disintegrated with accutase and incubated at 37°C for 30 min. Plates were washed and homogenized, then the cells were centrifuged, resuspended and cultured in stem cell medium. Flasks were incubated lying down to better select 3LL postC cells, since some cells remained attached to the surface and discarded in subsequent passages.

### Protein extraction for Immunoblotting

Cells were centrifuged and resuspended in extraction buffer, containing 1M Tris (pH 6.8), 10% SDS, 0.5M EDTA, 0.1M EGTA and supplemented with protease and phosphatase inhibitor cocktail set. The protein lysates were then heated, sonicated and centrifuged for 5 min each step. The Qubit kit was used to quantify proteins by fluorimetry and 20 ug of protein of each sample was diluted in RIPA buffer, containing 4x loading buffer and 10x reducing agent. Samples then were heated, centrifuged and incubated on ice for 5 min. Proteins were separated by denaturing electrophoresis using 4-12% Bis-Tris PAGE gels in 1X MES buffer. Gels run at 130V for 1h and then transferred to a nitrocellulose membrane using the iBlot2 Transfer Device (23V for 6 min). Membranes were blocked for 1h with 5% skimmed milk powder solution and incubated overnight at 4°C with primary antibodies (Table 1). The following day, membranes were washed 3 times and incubated with secondary antibodies (Table 1) at RT for 1h. Finally, membranes were incubated in ECL and the emitted light was detected by a ChemiDoc MP Imaging System transilluminator.

### Immunocytochemistry

Cells were adhered to round covers pre-treated with poly-L-lysine and fixed with 4% paraformaldehyde in 0.1M phosphate buffer. Then, cells were washed, covered with antibody solution (PBS containing lysine in 10% FBS) and stored at 4°C until further use. Finally, cells were incubated in blocking solution (0.2% Triton X-100 in antibody solution) in stirring for 1h and incubated with primary antibodies (Table 1) overnight at 4°C. Next day, cells were washed and incubated with secondary antibodies (Table 1) at RT and darkness for 1h. Nuclei were stained with DAPI for 5 min, then washed twice and the covers were mounted on slides in the presence of ProLong mountant medium.

### Image analysis

The intensity of the dots obtained with western blot was measured with the Fiji-ImageJ program. Immunostained cells were imaged with an Olympus AX70 fluorescence microscope (40X objective) and analyzed also with Fiji-ImageJ.

### Cell survival assays

Cells were directly seeded in 96-well plates (15×10^3^ cells per well in 200 μL) and treated with vehicle (DMSO) or increasing concentrations of the indicated inhibitors for 48 hours. Final concentration of DMSO was kept at 0.1%. Cells were incubated with 5 % resazurin for 4 hours, and fluorescence was measured at λ_ex_ = 565 nm and λ_em_ = 590 nm. Kinase inhibitor treatments were performed in three biological replicates. The list of kinase inhibitors used in this study is located in Table 2.

### Sample preparation for proteomic and phosphoproteomic analysis

After cell counting, 3LL-CSCs cells were seeded in T25 flasks (10^4^ cells/mL) in stem cell medium and maintained in an incubator overnight at 37°C and 5% CO_2_. Cells were harvested by centrifugation at 500 x *g* at 4°C for 5 min, washed twice with ice-cold PBS (supplemented with 1mM Na_3_VO_4_ and 1 mM NaF) and lysed in 500 μL urea buffer (8 M urea in 20 mM in HEPES pH 8.0 supplemented with 1 mM Na_3_VO_4_, 1 mM NaF, 1mM Na_4_P_2_O_7_ and 1 mM sodium β-glycerophosphate). All cell lysates were homogenized by sonication and insoluble material was removed by centrifugation at 16,000 x *g* for 15 min at 4°C. Protein in the cell extracts was quantified by bicinchoninic acid analysis.

For proteomics and phosphoproteome analysis, we used published methods [36–38], with some modifications. Briefly, 50 or 110 µg of protein were reduced and alkylated by sequential incubation with 10 mM DTT and 17 mM iodoacetamide for 1 hour. The urea concentration was diluted to 2 M with 20 mM HEPES (pH 8.0) and 100 µL of conditioned trypsin beads (50% slurry of TLCK-trypsin) (Thermo-Fisher Scientific; Cat. 20230) were added and the samples incubated for 16 hours at 37°C with agitation. Trypsin beads were removed by centrifugation at 2,000 x *g* for 5 min at 4°C. Samples were transferred to 96-well plates and subsequently acidified adding trifluoroacetic acid (TFA) to a final concentration of 0.1%. Desalting and phosphopeptide enrichment (this further step only for phosphoproteomics experiments) procedures were performed in the AssayMAP Bravo platform (Agilent Technologies) and the eluted samples were dried in a speedvac. Peptide pellets were stored at -80°C.

### MS-based proteomics and phosphoproteomics

Peptide and phosphopeptide pellets were re-suspended in reconstitution buffer (20 fmol/µL enolase in 3% ACN, 0.1% TFA) and loaded onto an LC-MS/MS system consisting of a Dionex UltiMate 3000 RSLC directly coupled to an Orbitrap Q-Exactive Plus mass spectrometer (Thermo Fisher Scientific). The LC system used mobile phases A (3% ACN; 0.1% FA) and B (100% ACN; 0.1% FA). Peptides were trapped in a μ-pre-column (cat. 160454) and separated in an analytical column (Acclaim PepMap 100, cat. 164569). The following parameters were used: 3% to 23% B gradient for 120 or 60 min, respectively and a flow rate of 0.3 µL/min.

As they eluted from the nano-LC system, peptides were infused into the online connected Q-Exactive Plus system operating with a 2.1s duty cycle. Acquisition of full scan survey spectra (m/z 375-1,500) with a 70,000 FWHM resolution was followed by, data-dependent acquisition in which the 15 most intense ions were selected for HCD (higher energy collisional dissociation) and MS/MS scanning (200-2,000 m/z) with a resolution of 17,500 FWHM. A 30 s dynamic exclusion period was enabled with an exclusion list with 10 ppm mass window. Overall duty cycle generated chromatographic peaks of approximately 30 s at the base, which allowed the construction of extracted ion chromatograms (XICs) with at least 10 data points.

### Peptide identification from tandem mass spectrometry data

Mascot Daemon (v2.8.0.1) was used to automate peptide identification from MS data. Peak list files from RAW data were generated with Mascot Distiller and loaded into the Mascot search engine in order to match MS/MS data to peptides [39]. The searches were performed against the SwissProt Database (SwissProt_2021_02. fasta) specifically for *Mus musculus*, with an FDR of ∼1% and the following parameters: 2 trypsin missed cleavages, mass tolerance of ±10 ppm for the MS scans and ±25 mmu for the MS/MS scans, carbamidomethyl Cys as a fixed modification, pyroGlu on N-terminal Gln and oxidation of Met as variable modifications. For phosphoproteomic experiments the phosphorylation on Ser, Thr and Tyr were included as variable modifications.

### Peptide quantification from MS1 data

The in-house developed Pescal software was used for label-free peptide quantification [40,41], XICs for all the peptides identified across all samples were constructed with ±7 ppm and ±2 min mass and retention time windows, respectively. Peak areas from all XICs were calculated. Undetectable peptides were given an intensity value of 0. Values of 2 technical replicates per sample were averaged and intensity values for each peptide were normalized to total sample intensity.

### Bioinformatics and statistical rationale

Pescal outputs were further processed with the protools2 v0.2.9 package (https://github.com/CutillasLab/protools2/). Protools2 read and normalized proteomics and phosphoproteomics data, performed statistical analysis using Limma [42] and Student t-test. Downstream processing was reimplemented in Python (v3.12) for deeper analysis, including KSEA and pathway enrichment analysis. Welch’s t-test was performed to assess significance in proteomics and phosphoproteomics experiments, two-way ANOVA and further Bonferroni’s or Dunnett’s multiple comparisons tests were used to assess significance in western blot and cell survival data. Where applicable, p-values were adjusted for multiple testing using Benjamini-Hochberg method

## Data and software availability

The mass spectrometry-based proteomics and phosphoproteomics data have been deposited to the ProteomeXchange Consortium via the PRIDE [43] partner repository with the dataset identifier PXD086427 and 10.6019/PXD086427.

## Author Contributions

E.A.R and P.C.G conducted experiments, A.F.F, A.M.C, B.F.R and P.R.C performed bioinformatics experiments, provided critical analysis and edited manuscript, V.R performed mass spectrometry experiments, A.T and R.C designed experiments, provided critical analysis and edited manuscript, M.H conceived study, designed and conducted experiments, analysed and interpreted data, prepared figures and wrote manuscript. All authors have read and agreed to the published version of the manuscript.

## Data Availability Statement

The original contributions presented in this study are included in the article. Further inquiries can be directed to the corresponding author.

## Acknowledgments

The authors specially thank Dr. Miguel Pericacho from University of Salamanca for kindly providing the 3LL parental cell line. The authors wish to extend their special thank to COST Action CA22103 Net4Brain (A Comprehensive Network Against Brain Cancer), supported by COST (European Cooperation in Science and Technology).

## Abbreviations

The following abbreviations are used in this article:

3LL postC: 3LL intracranially post-implanted cells
3LL preC: 3LL pre-implanted cells
ACN: acetonitrile
CNS: central nervous system
CSCs: cancer stem cells
DMSO: dimethyl sulfoxide
FA: formic acid
FDR: false discovery rate
ICC: immunocytochemistry
KSEA: kinase substrate enrichment analysis
MHC-I: major histocompatibility complex class I
MS: mass spectrometry
PBS: phosphate buffered saline
TFA: trifluoroacetic acid
XICs: extracted ion chromatograms

## Supplementary Tables

**Supplementary Table 1.**
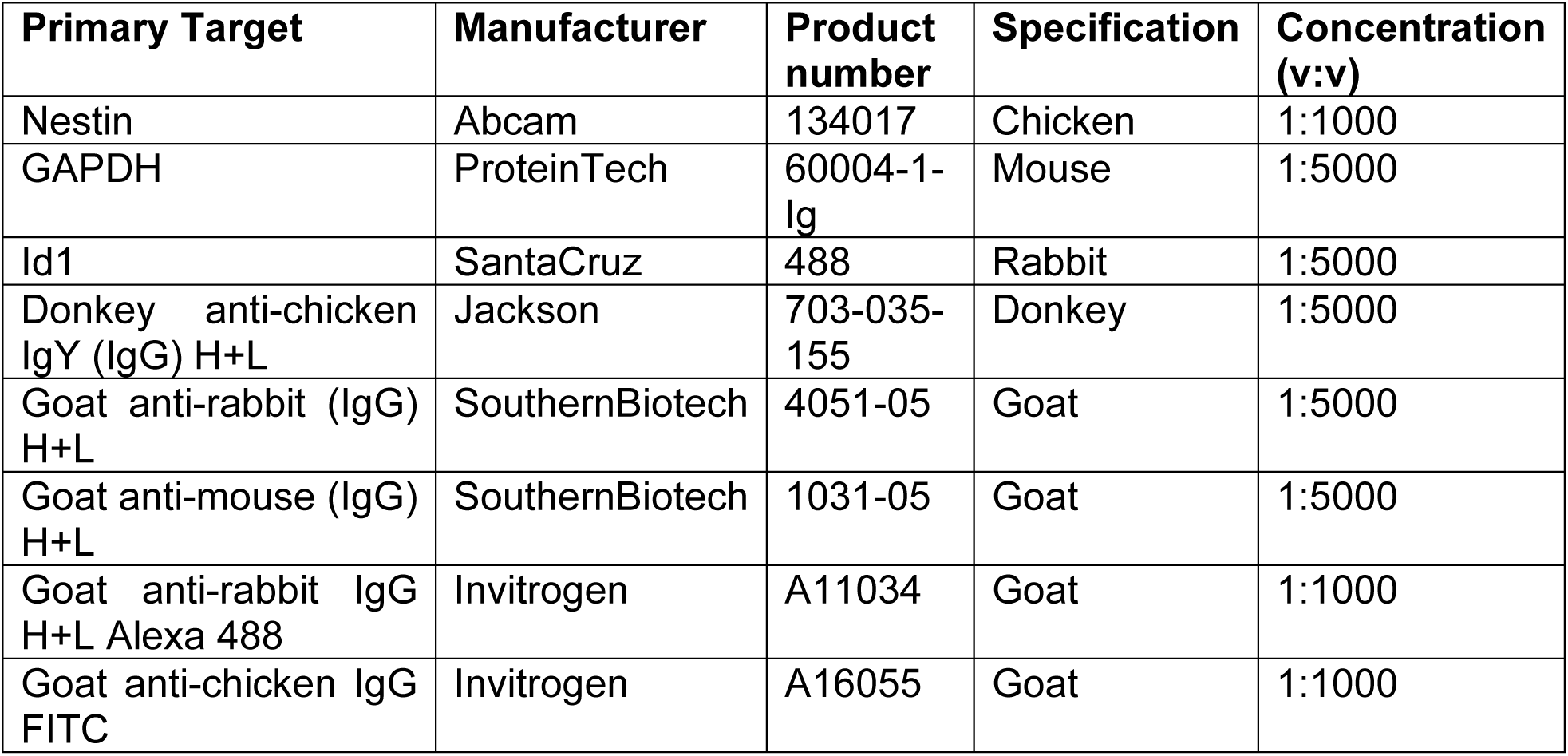
List of antibodies used for immunodetection.

**Supplementary Table 2.** List of kinase inhibitors used.

| Kinase inhibitor | Manufacturer | Catalog number | Primary target | IC50 (nM) |
| --- | --- | --- | --- | --- |
| PKC theta inhibitor | TargetMol | T5423 | PKCθ | 12 |
| Takinib | TargetMol | T4264 | TAK1 | 9.5 |
| Sotrastaurin | MedChemExpress | HY-10343 | pan-PKC | - |
| Darovasertib | MedChemExpress | HY-101569 | PKCα, PKCθ, GSK3β | 1.9 / 0.4 / 3.1 |

## References

1. Thai AA, Solomon BJ, Sequist LV, Gainor JF, Heist RS (2021) Lung cancer. Lancet 398 (10299):535–554. doi:10.1016/s0140-6736(21)00312-3

2. Boire A, Brastianos PK, Garzia L, Valiente M (2020) Brain metastasis. Nat Rev Cancer 20 (1):4–11. doi:10.1038/s41568-019-0220-y

3. Valiente M, Anders C, Boire A, Izar B, Kotecki N, Malladi S, Massagué J, Moss NS, Neman J, Preusser M, Schagen S, Siegel PM, Tawbi H, Venkataramani V, Winkler F, Zadeh G, Joyce JA (2026) The evolving landscape of brain metastasis: volume II. Trends Cancer 12 (2):116–146. doi:10.1016/j.trecan.2025.11.007

4. Suh JH, Kotecha R, Chao ST, Ahluwalia MS, Sahgal A, Chang EL (2020) Current approaches to the management of brain metastases. Nat Rev Clin Oncol 17 (5):279–299. doi:10.1038/s41571-019-0320-3

5. Quail DF, Joyce JA (2017) The Microenvironmental Landscape of Brain Tumors. Cancer Cell 31 (3):326–341. doi:10.1016/j.ccell.2017.02.009

6. Priego N, Valiente M (2019) The Potential of Astrocytes as Immune Modulators in Brain Tumors. Front Immunol 10:1314. doi:10.3389/fimmu.2019.01314

7. Batlle E, Clevers H (2017) Cancer stem cells revisited. Nat Med 23 (10):1124–1134. doi:10.1038/nm.4409

8. Lytle NK, Barber AG, Reya T (2018) Stem cell fate in cancer growth, progression and therapy resistance. Nat Rev Cancer 18 (11):669–680. doi:10.1038/s41568-018-0056-x

9. Er EE, Valiente M, Ganesh K, Zou Y, Agrawal S, Hu J, Griscom B, Rosenblum M, Boire A, Brogi E, Giancotti FG, Schachner M, Malladi S, Massagué J (2018) Pericyte-like spreading by disseminated cancer cells activates YAP and MRTF for metastatic colonization. Nat Cell Biol 20 (8):966–978. doi:10.1038/s41556-018-0138-8

10. Atanasescu VP, Breazu A, Oprea S, Porosnicu AL, Oproiu A, Rădoi MP, Munteanu O, Pantu C (2025) The Neuro-Melanoma Singularity: Convergent Evolution of Neural and Melanocytic Networks in Brain Metastatic Adaptation. Biomolecules 15 (12). doi:10.3390/biom15121683

11. Redmer T (2018) Deciphering mechanisms of brain metastasis in melanoma - the gist of the matter. Mol Cancer 17 (1):106. doi:10.1186/s12943-018-0854-5

12. Boiko AD, Razorenova OV, van de Rijn M, Swetter SM, Johnson DL, Ly DP, Butler PD, Yang GP, Joshua B, Kaplan MJ, Longaker MT, Weissman IL (2010) Human melanoma-initiating cells express neural crest nerve growth factor receptor CD271. Nature 466 (7302):133–137. doi:10.1038/nature09161

13. Cordero A, Kanojia D, Miska J, Panek WK, Xiao A, Han Y, Bonamici N, Zhou W, Xiao T, Wu M, Ahmed AU, Lesniak MS (2019) FABP7 is a key metabolic regulator in HER2+ breast cancer brain metastasis. Oncogene 38 (37):6445–6460. doi:10.1038/s41388-019-0893-4

14. Morgan R, Pandha HS (2020) PBX3 in Cancer. Cancers (Basel) 12 (2). doi:10.3390/cancers12020431

15. Yang L, Rau R, Goodell MA (2015) DNMT3A in haematological malignancies. Nat Rev Cancer 15 (3):152–165. doi:10.1038/nrc3895

16. Perissi V, Jepsen K, Glass CK, Rosenfeld MG (2010) Deconstructing repression: evolving models of co-repressor action. Nat Rev Genet 11 (2):109–123. doi:10.1038/nrg2736

17. Valiente M, Obenauf AC, Jin X, Chen Q, Zhang XH, Lee DJ, Chaft JE, Kris MG, Huse JT, Brogi E, Massagué J (2014) Serpins promote cancer cell survival and vascular co-option in brain metastasis. Cell 156 (5):1002–1016. doi:10.1016/j.cell.2014.01.040

18. Seano G, Nia HT, Emblem KE, Datta M, Ren J, Krishnan S, Kloepper J, Pinho MC, Ho WW, Ghosh M, Askoxylakis V, Ferraro GB, Riedemann L, Gerstner ER, Batchelor TT, Wen PY, Lin NU, Grodzinsky AJ, Fukumura D, Huang P, Baish JW, Padera TP, Munn LL, Jain RK (2019) Solid stress in brain tumours causes neuronal loss and neurological dysfunction and can be reversed by lithium. Nat Biomed Eng 3 (3):230–245. doi:10.1038/s41551-018-0334-7

19. Kienast Y, von Baumgarten L, Fuhrmann M, Klinkert WE, Goldbrunner R, Herms J, Winkler F (2010) Real-time imaging reveals the single steps of brain metastasis formation. Nat Med 16 (1):116–122. doi:10.1038/nm.2072

20. Lendahl U, Zimmerman LB, McKay RD (1990) CNS stem cells express a new class of intermediate filament protein. Cell 60 (4):585–595. doi:10.1016/0092-8674(90)90662-x

21. Klemm F, Maas RR, Bowman RL, Kornete M, Soukup K, Nassiri S, Brouland JP, Iacobuzio-Donahue CA, Brennan C, Tabar V, Gutin PH, Daniel RT, Hegi ME, Joyce JA (2020) Interrogation of the Microenvironmental Landscape in Brain Tumors Reveals Disease-Specific Alterations of Immune Cells. Cell 181 (7):1643–1660.e1617. doi:10.1016/j.cell.2020.05.007

22. Sevenich L (2018) Brain-Resident Microglia and Blood-Borne Macrophages Orchestrate Central Nervous System Inflammation in Neurodegenerative Disorders and Brain Cancer. Front Immunol 9:697. doi:10.3389/fimmu.2018.00697

23. Casado P, Rodriguez-Prados JC, Cosulich SC, Guichard S, Vanhaesebroeck B, Joel S, Cutillas PR (2013) Kinase-substrate enrichment analysis provides insights into the heterogeneity of signaling pathway activation in leukemia cells. Sci Signal 6 (268):rs6. doi:10.1126/scisignal.2003573

24. Altman A, Villalba M (2003) Protein kinase C-theta (PKCtheta): it’s all about location, location, location. Immunol Rev 192:53–63. doi:10.1034/j.1600-065x.2003.00027.x

25. Mihaly SR, Ninomiya-Tsuji J, Morioka S (2014) TAK1 control of cell death. Cell Death Differ 21 (11):1667–1676. doi:10.1038/cdd.2014.123

26. Perk J, Iavarone A, Benezra R (2005) Id family of helix-loop-helix proteins in cancer. Nat Rev Cancer 5 (8):603–614. doi:10.1038/nrc1673

27. Wang R, Khan S, Liao G, Wu Y, Tang DD (2022) Nestin Modulates Airway Smooth Muscle Cell Migration by Affecting Spatial Rearrangement of Vimentin Network and Focal Adhesion Assembly. Cells 11 (19). doi:10.3390/cells11193047

28. Ruzinova MB, Benezra R (2003) Id proteins in development, cell cycle and cancer. Trends Cell Biol 13 (8):410–418. doi:10.1016/s0962-8924(03)00147-8

29. Redmer T, Welte Y, Behrens D, Fichtner I, Przybilla D, Wruck W, Yaspo ML, Lehrach H, Schafer R, Regenbrecht CR (2014) The nerve growth factor receptor CD271 is crucial to maintain tumorigenicity and stem-like properties of melanoma cells. PLoS One 9 (5):e92596. doi:10.1371/journal.pone.0092596

30. Thakker-Varia S, Behnke J, Doobin D, Dalal V, Thakkar K, Khadim F, Wilson E, Palmieri A, Antila H, Rantamaki T, Alder J (2014) VGF (TLQP-62)-induced neurogenesis targets early phase neural progenitor cells in the adult hippocampus and requires glutamate and BDNF signaling. Stem Cell Res 12 (3):762–777. doi:10.1016/j.scr.2014.03.005

31. Infante P, Faedda R, Bernardi F, Bufalieri F, Lospinoso Severini L, Alfonsi R, Mazzà D, Siler M, Coni S, Po A, Petroni M, Ferretti E, Mori M, De Smaele E, Canettieri G, Capalbo C, Maroder M, Screpanti I, Kool M, Pfister SM, Guardavaccaro D, Gulino A, Di Marcotullio L (2018) Itch/β-arrestin2-dependent non-proteolytic ubiquitylation of SuFu controls Hedgehog signalling and medulloblastoma tumorigenesis. Nat Commun 9 (1):976. doi:10.1038/s41467-018-03339-0

32. Khazaei MR, Girouard MP, Alchini R, Ong Tone S, Shimada T, Bechstedt S, Cowan M, Guillet D, Wiseman PW, Brouhard G, Cloutier JF, Fournier AE (2014) Collapsin response mediator protein 4 regulates growth cone dynamics through the actin and microtubule cytoskeleton. J Biol Chem 289 (43):30133–30143. doi:10.1074/jbc.M114.570440

33. Omary MB, Ku NO, Tao GZ, Toivola DM, Liao J (2006) “Heads and tails” of intermediate filament phosphorylation: multiple sites and functional insights. Trends Biochem Sci 31 (7):383–394. doi:10.1016/j.tibs.2006.05.008

34. Yi L, Zhou C, Wang B, Chen T, Xu M, Xu L, Feng H (2013) Implantation of GL261 neurospheres into C57/BL6 mice: a more reliable syngeneic graft model for research on glioma-initiating cells. Int J Oncol 43 (2):477–484. doi:10.3892/ijo.2013.1962

35. Jaraíz-Rodríguez M, Talaverón R, García-Vicente L, Pelaz SG, Domínguez-Prieto M, Álvarez-Vázquez A, Flores-Hernández R, Sin WC, Bechberger J, Medina JM, Naus CC, Tabernero A (2020) Connexin43 peptide, TAT-Cx43266-283, selectively targets glioma cells, impairs malignant growth, and enhances survival in mouse models in vivo. Neuro Oncol 22 (4):493–504. doi:10.1093/neuonc/noz243

36. Hijazi M, Smith R, Rajeeve V, Bessant C, Cutillas PR (2020) Reconstructing kinase network topologies from phosphoproteomics data reveals cancer-associated rewiring. Nat Biotechnol 38 (4):493–502. doi:10.1038/s41587-019-0391-9

37. Hijazi M, Casado P, Akhtar N, Alvarez-Teijeiro S, Rajeeve V, Cutillas PR (2022) eEF2K Activity Determines Synergy to Cotreatment of Cancer Cells With PI3K and MEK Inhibitors. Mol Cell Proteomics 21 (6):100240. doi:10.1016/j.mcpro.2022.100240

38. Cerveró-García P, Jiménez-Madrona E, Álvarez-Vázquez A, Flores-Hernández R, García-Vicente L, García-Guerrero L, Alonso-Amador JJ, González-Sánchez R, Ollauri-Ibáñez C, Paniagua-Sancho M, Talaverón R, Rodrigues-Teixeira T, Martín-Guerrero SM, Casado P, Rajeeve V, Shields T, Hijazi M, Cutillas PR, Jaraíz-Rodríguez M, Tabernero A (2026) The Src inhibitor peptide TAT-Cx43. Neurooncol Adv 8 (1):vdag001. doi:10.1093/noajnl/vdag001

39. Perkins DN, Pappin DJ, Creasy DM, Cottrell JS (1999) Probability-based protein identification by searching sequence databases using mass spectrometry data. Electrophoresis 20 (18):3551–3567. doi:10.1002/(sici)1522-2683(19991201)20:183551<::Aid-elps3551>3.0.Co;2-2

40. Wilkes E, Cutillas PR (2017) Label-Free Phosphoproteomic Approach for Kinase Signaling Analysis. Methods Mol Biol 1636:199–217. doi:10.1007/978-1-4939-7154-1_13

41. Cutillas PR, Vanhaesebroeck B (2007) Quantitative profile of five murine core proteomes using label-free functional proteomics. Mol Cell Proteomics 6 (9):1560–1573. doi:10.1074/mcp.M700037-MCP200

42. Ritchie ME, Phipson B, Wu D, Hu Y, Law CW, Shi W, Smyth GK (2015) limma powers differential expression analyses for RNA-sequencing and microarray studies. Nucleic Acids Res 43 (7):e47. doi:10.1093/nar/gkv007

43. Perez-Riverol Y, Bai J, Bandla C, García-Seisdedos D, Hewapathirana S, Kamatchinathan S, Kundu DJ, Prakash A, Frericks-Zipper A, Eisenacher M, Walzer M, Wang S, Brazma A, Vizcaíno JA (2022) The PRIDE database resources in 2022: a hub for mass spectrometry-based proteomics evidences. Nucleic Acids Res 50 (D1):D543–D552. doi:10.1093/nar/gkab1038

